# Apoplast multi-omics profiling during fungal infection uncovers new players of basal and early induced immunity

**DOI:** 10.64898/2026.01.28.702280

**Authors:** Antoine Davière, Sylvie Jolivet, Annie Auger, Gilles Clement, Stephanie Boutet, Jean Chrisologue Totozafy, Thierry Balliau, Mélisande Blein-Nicolas, Justine Rouffet, Marie-Christine Soulié, Mathilde Fagard

**Affiliations:** Université Paris-Saclay, INRAE, AgroParisTech, Institute Jean-Pierre Bourgin for Plant Sciences, 78000, Versailles, France; Université Paris Saclay, ED 567 Sciences du végétal, 91405 Orsay, France; Université Paris-Saclay, INRAE, CNRS, AgroParisTech, GQE-Le Moulon, PAPPSO, 91190, Gif-Sur-Yvette, France; Université Paris-Saclay, INRAE, AgroParisTech, BIOGER, 91405, Orsay, France; Sorbonne Université, Pierre et Marie Curie Université Paris 06, UFR 927, 75005 Paris, France

## Abstract

The extracellular space in plant tissues, known as the apoplast, remains one of the least characterized cellular compartments. The apoplastic fluid (APF), akin to mammalian extracellular fluid, serves as the primary interface between pathogens and their host. It allows the exchange of signalling molecules, coordinates host cell responses, and enables the circulation of pathogen effectors that modulate the immune response. We describe here the first multi-omics analysis of the APF content just 6 h after the onset of *A. thaliana* infection with *B. cinerea*, a fast-killing necrotrophic fungus. By varying plant nitrogen nutrition, known to affect both plant defenses and pathogen virulence, we identify candidates that do not stand out under optimal conditions. Our analysis uncovers novel nitrogen-dependent mechanisms that regulate both basal and early induced apoplastic immunity, revealing the presence of previously unidentified, potentially protective apoplastic metabolites as well as the intercellular transport of nuclear proteins, thereby offering new insights into early apoplastic immune responses.

## Introduction

All organisms are subjected during their lifetime to different stresses, of abiotic and biotic origins, imposed by the environment. Whether plants or animals—hosts often employ shared defensive strategies to repel pathogens, such as physical barriers and innate immunity. In turn, parasites have developed parallel tactics to overcome these defenses, using methods like active invasion and evasion of the host’s innate and adaptive immune systems^1^. To protect themselves from pathogen infection, plants possess different layers of defense, the first being non-specific preformed barriers that include the cuticle and the cell wall as well as chemicals such as phytoanticipins that efficiently restrain some pathogens. In addition, plants can induce specific defenses upon the recognition of microbial elicitors such as pathogen-or damage-associated molecular patterns (PAMPs and DAMPs). Their recognition by specific receptors activates PAMP-triggered immunity (PTI) which rapidly induces a strong reactive oxygen species (ROS) production both for signalling and direct defensce purposes. Conjointly, co-evolution of plants and pathogens has led to the sophisticated recognition of pathogen effectors through specific defense proteins through a mechanism called effector triggered immunity (ETI). In reaction to these perception mechanisms, plant hormones like jasmonate (JA), salycilate (SA) and ethylene (ET) are produced, specifically orchestrating the defense reaction, as highlighted by their pathogen-type dependent level of accumulation^2^. Altogether this leads to the induction of defense-associated gene expression, remodelling of the cell wall, production of defense-related primary and specialized metabolites^3^ and accumulation of defense proteins such as the pathogenesis-related (PR) proteins in the extracellular apoplastic space.

The apoplastic space, defined as the extracellular matrix including the cell wall and the intercellular space containing the apoplastic fluid (APF), is the first plant compartment where pathogen-host interactions occur^4^. The APF shares similarities with extracellular fluids in mammals, both containing a specific array of molecules that regulate key physiological functions, such as development, response to abiotic stress and pathogen defense^1^. It allows the exchange of signalling molecules between plant cells and the coordination of their response following pathogen perception. Recent advances in scRNAseq, spatial transcriptomic and epigenomic enabled the discovery of two kinds of early activated immune cell states, PRIMER cells, at the pathogen contact site, and bystander cells that are in the surroundings^5^. Cell-cell communication through the apoplastic and symplastic routes between the co-existent specific immune cell states likely plays an important role coordinating the response but is still poorly characterized^6^. In addition to cell-cell communication within the host, proteins and metabolites circulating in the APF contribute to the early molecular dialogue between host cells and pathogens, conditioning the outcome of the interaction. A tight control of available nutrients like sugars and amino acids in the apoplast around the infection site is critical for both pathogen nutrition and immune signalling^7^. For example, inactivating or ectopically overexpressing sugar or amino acid transporters in plants was shown to increase resistance through pathogen starvation or nutrient-triggered immunity mechanisms^8,9^. Another important process taking place in the APF is the secretion and activation of host cell wall-remodelling enzymes that not only shape the physical barrier and limit degradation by pathogen cell wall-degrading enzymes (CWDEs) but also regulate cell wall-derived signalling. Recent works have investigated the complexity of oligogalacturonides (OGs) signalling in the apoplast^10–13^. In addition to direct secretion, both plant and mammalian organisms use extracellular vesicles (EVs) to deliver molecules to the extracellular space and beyond^1^. Recent characterization of these EVs revealed their prominent role in host-pathogen interactions, in plants and in mammals. Produced from both the pathogen and the host side, these vesicles can carry siRNAs to repress pathogen virulence or plant defense^14–16^. More strikingly, *Arabidopsis thaliana* mRNA detected in EVs can be delivered in fungal cells and translated into proteins repressing virulence^17^. A similar mechanism probably exists from the pathogen side but still needs to be validated through the challenging detection of low abundant pathogen-derived EVs.

Important modifications in the proteome and/or the metabolome of the APF have been described following perception of different pathogens or elicitors^18–27^. These studies reported the accumulation of well-known defense proteins such as peroxidases, CWDEs and proteases as well as hormones, primary and specialized metabolites in this compartment. More recently, several studies have demonstrated that characterizing the apoplastic fluid (APF) provides critical insights into pathogen virulence *in planta*. For example, characterization of non-infected APF extracts with transmission electronic microscopy and proteomics revealed the presence of host proteasome subunits^28^. Using the proteasome inhibitor syringolin-A produced by the bacterial pathogen *Pseudomonas syringae*, the authors showed that the extracellular proteasome is involved in processing flagellin into its immune peptide flg22 enabling PTI induction. In addition, our previous work demonstrated that in *Arabidopsis thaliana* apoplastic fluid (APF), specific metabolites which induce *Erwinia amylovora* virulence genes accumulate differentially depending on plant nitrogen (N) supply—directly correlating with enhanced bacterial fitness^21^. Thus, while providing information on host defense strategies taking place in the APF to limit pathogen development, studying this compartment can also reveal new secreted pathogenicity factors as well as plant signals required to activate them. Indeed, only a few studies have successfully detected pathogen proteins in the APF and characterized their role as effectors^24,25^. Most of these studies, in particular omics-based analysis, have focused on biotrophic and hemi-biotrophic fungi or apoplastic-dividing bacteria, whose lifestyle, contrary to necrotrophs, involves prolonged molecular dialogue with the host in the apoplast.

The broad host-range fungal pathogen *Botrytis cinerea*, an ascomycete of the *Sclerotiniaceae* family, is known to secrete a large diversity of effectors in the APF to modulate plant immunity towards the induction of plant-cell death which is favourable for its necrotrophic lifestyle^29^. Several such *B. cinerea* effectors, collectively termed cell death-inducting proteins (CDIPs), were previously characterized such as BcIEB1, BcSPL1, BcXIG1, BcXYN11a and BcCRH1^30–34^. Nevertheless, Leisen et al (2022) revealed through the production of multiple knockout mutants, the high level of redundancy of these effectors. Indeed, depending on the host plant tested, the deletion of up to 12 CDIP-coding genes is necessary to significantly reduce pathogenicity. In addition, *B. cinerea* produces many other different types of pathogenicity factors such as proteases, ROS-detoxifying enzymes, toxins and CWDEs. For example, the abundant and early secreted polygalacturonase BcPG1 but also the pectin-lyase BcPNL1 highly contribute to pathogenicity^10,36^. It is now well established that the initial phase of the infection process (before 24 hour post inoculation) is critical and defines the fate of *B. cinerea* interaction with its hosts^37^. Accordingly, joint efforts in the field are done to uncover the early events of the interaction.

To obtain a unique insight into the apoplastic dialogue of a fast-killing necrotrophic fungus with its host we describe here the leaf apoplastic fluid content just 6 h after the onset of *A. thaliana* infection with *B. cinerea*. To identify factors that do not stand out under optimal conditions, we added a comparison with plants grown in N-limiting conditions that present more resistance to the fungus^10,38^. Collectively, this study provides a leaf APF multi-omics description of the interaction between plant N nutrition and pathogen infection and uncovers new players of basal and early induced apoplastic immunity as well as new putative pathogen effectors involved in the *B. cinerea*-*A. thaliana* interaction.

## Results

### Apoplast washing fluids (AWF) extraction from infected leaves

To investigate the leaf apoplastic space during the *A. thaliana* – *B. cinerea* interaction we first selected suitable conditions for the recovery of apoplast washing fluids (AWF) extracts (Figure 1). To perform the infection, the aerial part of 6-week-old plants was harvested and full-grown mature leaves were inoculated with large mycelial plugs to maximize the surface of contact sites between the mycelium and the leaf epidermis. As shown in Figure 1C, this method led to the production of the typical maceration symptoms of the grey mold disease 24 hours after inoculation (24 hpi). Then, we determined an optimal early observation time point that allowed detection of leaf responses to the fungus while minimizing tissue degradation and cytoplasmic leakage into the apoplast. To monitor tissue degradation, we performed a 3, 3’-diaminobenzidine (DAB) staining, which reveals the presence of H_2_O_2_, as well as a trypan blue (TB) staining which stains dying cells. At 24 hpi, strong TB and DAB stainings were visible, revealing respectively extensive cell death and oxidative burst (Figure 1D). Conversely, leaves sampled at 6 hpi presented no visible symptoms (Figure 1A) and TB staining at this time point revealed only small coloration spots coming in part from dying plant cells but also from staining of the mycelium itself (Figure 1B). However, at 6 hpi we observed large areas of DAB-staining, revealing H_2_O_2_ production at different sites of the leaves and confirming an extensive reaction of the leaf to the fungal infection at this early time point (Figure 1B). We thus selected this early time-point (6 hpi), that we previously explored with an RNAseq experiment^38^, for AWF extraction from the leaves.

**Figure 1:**
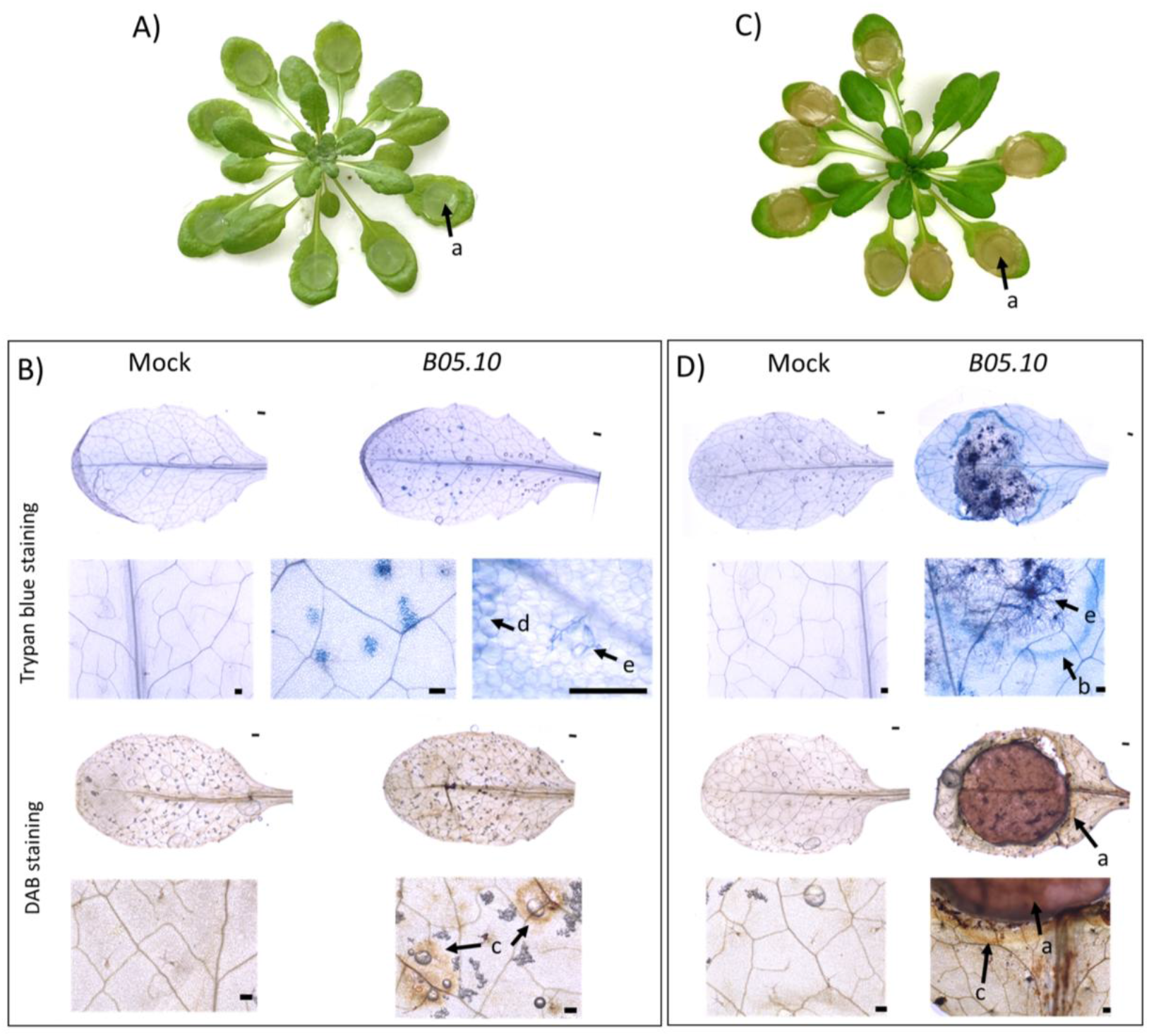
Detection of plant cellular reactions to fungal infection before the onset of maceration symptoms. (A, C) Representative picture of a 6-week-old A. thaliana plant 6 (A) or 24 (C) hours post inoculation with large mycelial plugs of B. cinerea. (B, D) A. thaliana leaves were tested for cell death reaction with trypan blue and for H_2_O_2_ production with 3, 3’-diaminobenzidine (DAB) at 6 hpi (B) and 24 hpi (D) with mock treatment or mycelial explants of the B. cinerea B05.10 wild-type (WT) strain. Arrows indicate the mycelial explant (a), extensive cell death (b), oxidative burst reaction (c), localized necrosis (d), infecting mycelium (e). Pictures were taken with the ZEISS Axio Zoom V16. About 4 leaves of independent plants per condition were observed at each time point and the experiment was repeated 3 times. Scale bar = 200µm.

The AWF composition is highly dependent on the infiltration solution used^39^. In this study we used ddH2O (double distilled water) as infiltration solution for extracts designated for metabolomic analysis, as in our earlier work^21^, to prevent any influence of buffer molecules on the detection of endogenous metabolites. Extracts for the proteomic analysis were obtained with an infiltration solution of mannitol and CaCl2 to increase the recovery of proteins by solubilization of weakly bound cell wall proteins^40^. Following infiltration of the harvested leaves, a low g centrifugation was applied to the leaves to recover the AWF^22^. The obtained AWF extracts were then tested for cytoplasmic contamination using activity of the cytoplasmic enzyme malate dehydrogenase (MDH) and total protein concentration as readouts. We observed no significant differences in MDH activity between mock AWF extracts (mAWF) and infected AWF extracts (iAWF) with both infiltration solutions (Table S1). Furthermore, the MDH activity levels observed in our study were similar to those obtained in other studies^22^. We observed a slight increase of total protein concentration in iAWF as it was observed in ^23^ and a higher concentration of proteins with the Mannitol-CaCl_2_ infiltration technique. Finally, we analysed the dilution factor of the AWF extracts as described previously^21,41^. The dilution factors were very similar between mock and infected sample with an average dilution factor of 1.67 and 1.65, respectively (Table S1) indicating no effect of the infection on AWF dilution at 6 hpi.

Altogether these results showed that our experimental approach enabled the collection of comparable mAWF and iAWF extracts with low cytoplasmic contaminations.

### Antifungal properties of AWF extracts

To determine whether our AWF extracts contained antifungal properties we tested their effect on *B. cinerea* growth *in vitro*. By monitoring fungal growth *in vitro* during 48 h with laser nephelometry as described in Joubert et al (2010), we observed a significant reduction of fungal growth in the presence of a 1/10 dilution of both mAWF and iAWF extracts in PDB compared to the control (PDB) (Figure 2A). Interestingly, no significant differences were found in the growth inhibiting capacity of mAWF and iAWF. This result indicates the presence of molecules able to restrain fungal growth in the apoplastic space even before any pathogen attack. To better characterize the effect of AWF on the fungus we recovered the mycelium after 48 h of growth with the extracts and subjected it to DAB staining to visualize H_2_O_2_ production. The darker DAB staining observed with the mycelium treated with mAWF and iAWF extracts compared to the PDB control condition indicated the presence of a higher H_2_O_2_ concentration in the mycelium (Figure 2B). Interestingly, iAWF induced stronger fungal H_2_O_2_ production than mAWF, indicating a difference in ROS-triggering activity of these extracts.

**Figure 2:**
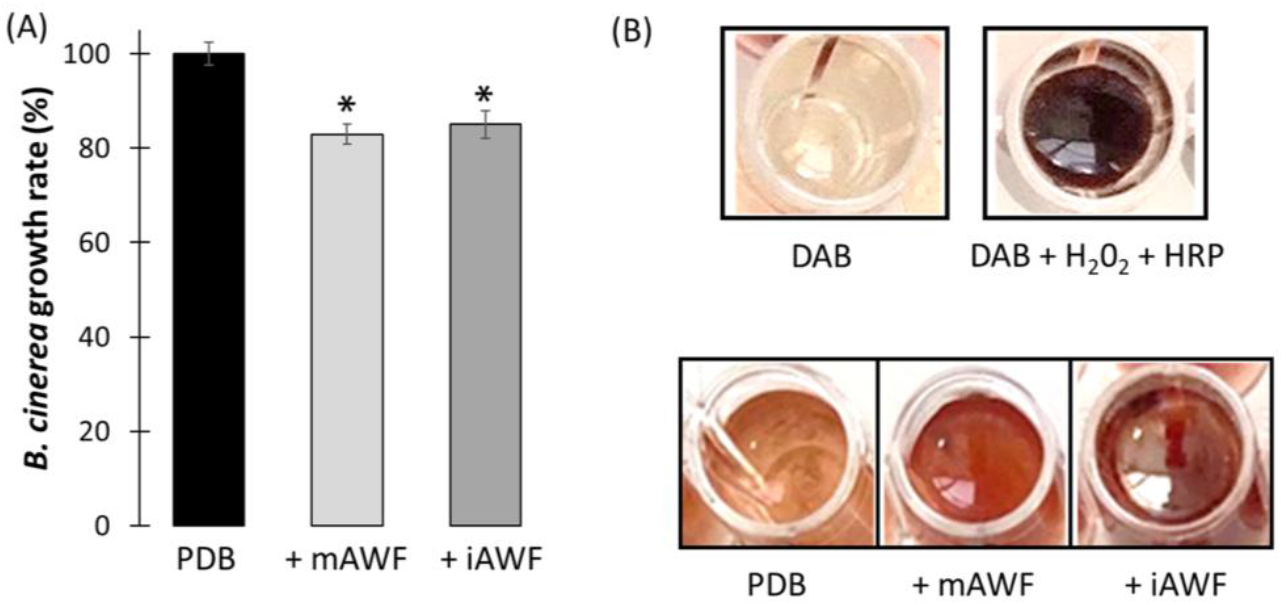
Antifungal effect of AWF extracts. (A) Spores of B. cinerea were grown in PDB with 1/10 dilution of different ddH_2_O AWF extracts (mAWF, iAWF) in 96-wells microplates. Growth represented by the OD_635nm_ was monitored every 15 min for 48 h in a nephelometric reader. The average growth rates were calculated during the exponential phase. Results are represented as the mean percentage of growth compared to the control condition (PDB, 100%). Experiments were repeated three times with four different biological AWF replicates. A two-sample t-test was performed (*p<0.05). (B) H_2_O_2_ production by B. cinerea following growth in mAWF and iAWF extracts compared to control conditions (PDB). At the end of the growth inhibition assay, each well of the microplate, was subjected to DAB staining to reveal H_2_O_2_ production. Two staining controls without the fungus were performed with either PDB + DAB, as a negative control, or PDB + DAB + 5 mM H_2_O_2_ + HRP (horseradish peroxidase) as a positive control. Experiments were repeated twice, and the same results were observed with four different biological replicas of AWF extracts.

### Primary metabolism shift toward defense in the apoplastic space following B. cinerea infection

Bacterial leaf infection has been shown previously to affect apoplastic primary metabolite content ^21,22^. Only a few studies have analysed fungal colonization on whole metabolomic apoplastic content and none concerned a necrotrophic fungus^23,43^. To assess whether early fungal infection with a necrotroph also led to primary metabolite changes in the apoplast, we performed GC-MS on mAWF and iAWF extracts. A total of 243 features were detected, of which 110 were annotated (Figure 3A). The most abundant metabolites found in mAWF were organic acids (fumarate, citrate, malate) followed by two sugars (fructose and glucose) and four amino acids (glutamate, aspartate, serine and GABA) (Figure 3B). Thirty one annotated features presented significant abundance variations, with 14 features up-accumulated in iAWF and 17 down-accumulated. Those 31 features were grouped into five categories: energy metabolism and amino acids, redox-related compounds, defense-related compounds and cell wall-derived compounds (Figure 3C). Concerning the energy metabolism category, metabolites were mostly depleted in iAWF compared to mAWF. Regarding redox-related compounds we found that in iAWF, dehydroascorbate (DHA) and ascorbate (AA), known to form an important redox couple regulating oxidative reactions, were both strongly depleted (Figure 3C). Interestingly, threonate which can originate from the catabolism of DHA and AA accumulated in iAWF. We also observed a depletion of phytol-2 in iAWF, a precursor of the antioxidant α-tocopherol. In addition, the oxidizable polyamine putrescine was accumulated in iAWF compared to mAWF.

**Figure 3:**
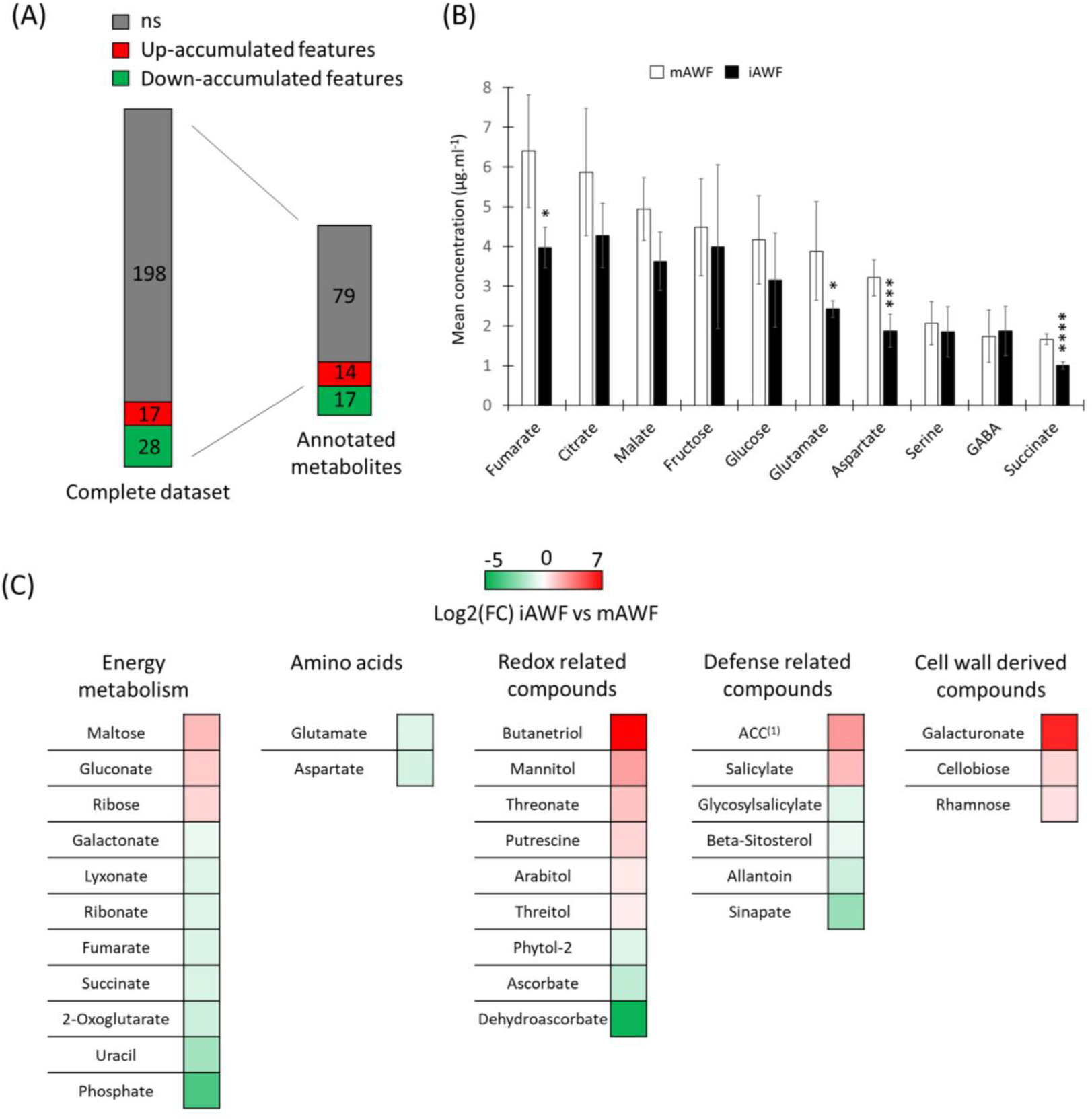
Early changes in apoplastic primary metabolites identified by GC-MS following A. thaliana infection with B. cinerea. AWF extracts were recovered 6 h following mock inoculation (mAWF) or inoculation with B. cinerea mycelial plugs (iAWF) and analyzed with GC-MS. (A) Schematic representation of the modifications in iAWF compared to mAWF in the complete dataset or only the annotated metabolites. Differently accumulated features were obtained with a paired t-test (p<0.05). ns = not significant. (B) The 10 most abundant primary metabolites recovered in AWFs extracts. Data represented are the mean of five biological replicates +/-SD. Paired t-test were done between mock AWF extracts and infected AWF extracts: *p<0.05, ***p<0.005, ****p<0.001. (C) Log2 fold changes of iAWF compared to mAWF are represented for annotated metabolite significantly different between the two conditions. Significant metabolites were identified with a paired t-test (p<0.05). Five biological replicates were analyzed to obtain these data. ^(1)^1-aminocyclopropane-1-carboxylate.

Concerning defense related compounds, we observed an accumulation of 1-aminocyclopropane-1-carboxylate (ACC) which is the precursor of ethylene (ET) and salicylate (SA) in iAWF whereas glycosylsalicylate, which is the inactive form of SA was depleted (Figure 3C). This result is consistent with the involvement of hormonal defense pathways in defense against *B. cinerea* infection^44^. Interestingly, a depletion of allantoin, which is a purine catabolite that can modulate JA-mediated defenses, was also observed in iAWF compared to mAWF^45–47^. Finally, consistent with the strong cell wall degrading capacities of *B. cinerea* we observed an accumulation, in iAWF compared to mAWF, of building blocks of cellulose and pectin i.e. cellobiose, galacturonate and rhamnose respectively.

To conclude, fungal infection led to a depletion in the apoplast of several energy-related metabolites whereas metabolites related to defense and the redox process accumulated in the apoplast following fungal infection. Moreover, the accumulation of cell wall-derived compounds and phytohormones in the apoplast after infection emphasizes their important role in apoplastic signalling and defense.

### Infection and nitrogen-dependent modulation of the AWF specialized metabolite content

Since the GC-MS analysis of AWF extracts described above unveiled a shift in primary metabolism towards building a defense response, we conducted an in-depth analysis of specialized metabolites in our extracts using untargeted LC-MS/MS. For this analysis, in addition to the extracts analysed above obtained under nitrogen (N)-sufficient conditions (10 mM NO_3_^-^, high N), we included extracts obtained from plants grown under low N (0.5 mM NO_3_^-^, low N). Indeed, *A. thaliana* plants grown under low N were previously shown to be more resistant to *B. cinerea* infection^10,38^. Accordingly, our aim was to use these infection conditions to reveal important apoplastic metabolites involved in plant resistance.

A total of 362 features were detected in the AWF extracts. By first selecting features significantly affected by the infection and/or by the combination of infection and plant N supply, we obtained a list of 20 metabolites grouped in five pathways: alkaloids, fatty acids, organosulfur, shikimates and phenylpropanoids and finally amino acids and peptides (Figure 4 A-E). Well-known defense metabolites such as alkaloids, including the camalexin toxin, and different types of octadecanoids, including the JA precursor OPDA, were strongly accumulated in iAWF compared to mAWF (Figure 4A, B). Five metabolites from various classes were significantly affected by the combination of infection and N nutrition (two-way ANOVA, inter, p<0.05). Unk#8, part of the fatty acids and conjugates class, was more accumulated in mAWF low N compared to high N but then depleted following infection in low N conditions (Figure 4B). This could indicate that this compound acts as a precursor for the synthesis of another molecule following infection in low N. The isothiocyanates 1-isothiocyanato-8-methylsulfinyl (8-MITC) and 1-isothiocyanato-7-methylsulfinyl (7-MITC), the flavonoid saponarin and the phenylpropanoid phenylacetaldehyde were all more accumulated in mAWF high N compared to all the other conditions that had a similar level (Figure 4C, D). These results indicate that unk#8 could be associated to basal resistance as a precursor for a defense compound produced in iAWF low N but not the four other metabolites. Next to these five metabolites, unk#11 a feature of m/z 663.1209 in ESI positive mode that was detected in very low amount in mAWF was strongly accumulated in iAWF of plants grown under low N compared to high N (Figure 4D). Although this tendency was not significant (two-way ANOVA, inter, p>0.05), the production of this compound could underlie the improved induced resistance of plants grown under low N. Using the fragmentation spectra of different detected adduct of this metabolite, we annotated this compound as a flavonoid with a raw formula C_29_HM_28_O_18_. The spectra used for the annotation and a putative structure of this flavonoid are shown in Figure S1.

**Figure 4:**
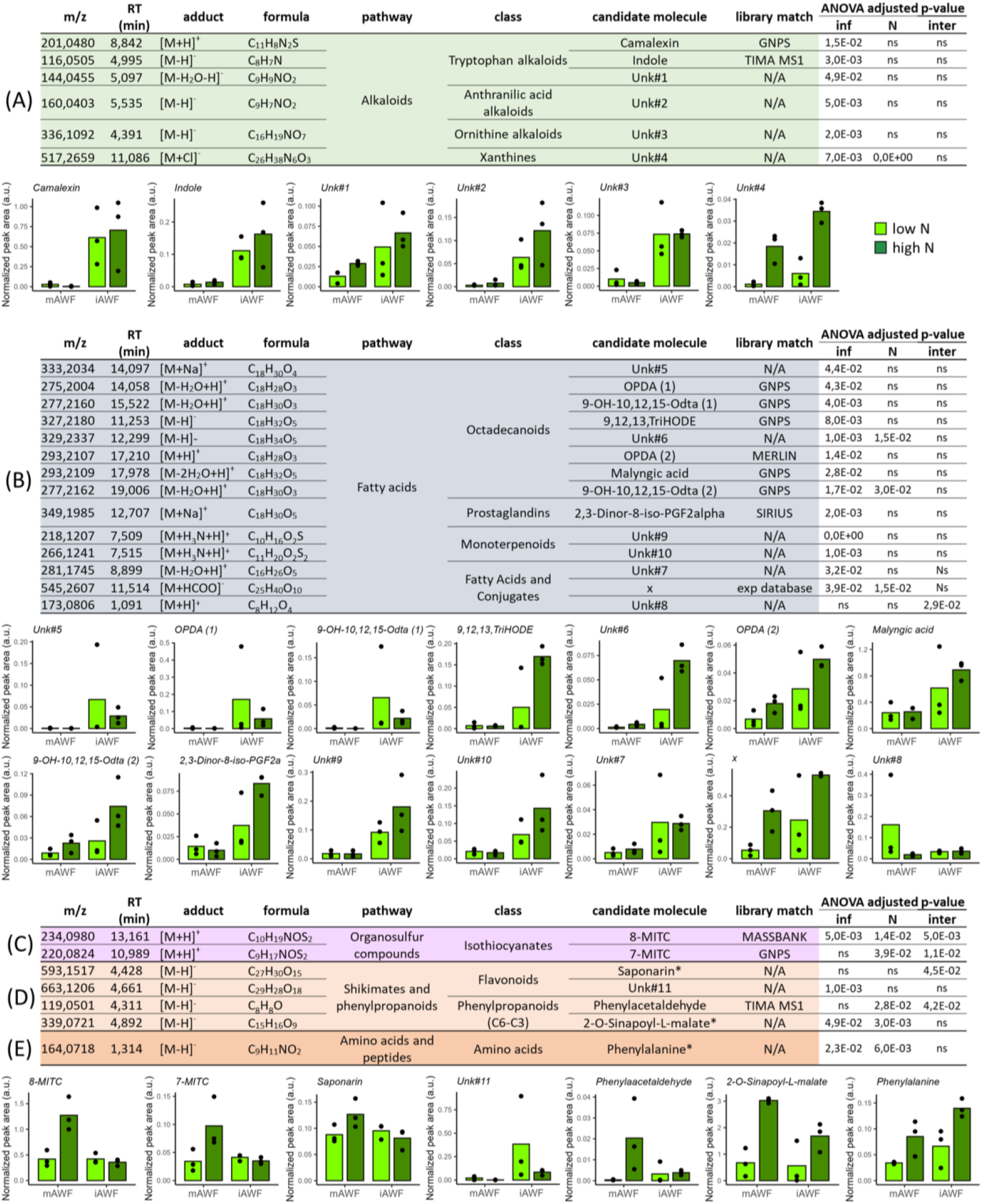
Early changes in apoplastic specialized metabolites following B. cinerea infection of A. thaliana plants grown with high or low nitrogen nutrition. AWF extracts from plants grown in high nitrogen (high N) or low nitrogen (low N) were recovered at 6 h post inoculation with mock (mAWF) or B. cinerea mycelial plugs (iAWF) before analysis with untargeted LC-MS/MS. (A, B, C, D, E) Tables show for each identified feature the detection parameters, the annotation details and the results of a two-way ANOVA for the two factors (inf, N) and their interaction (inter). Bar plots represent the mean of normalized peak area (a. u.) for each feature in mAWF and iAWF in low N or high N conditions with each dot representing one independent biological replicate. *: annotation validated by standards. Inf: infection status, N: N regime, inter: interaction. ^x^1-O-[2-Hydroxy-3-[6-[2-(2-pentenyl)-3-oxo-4-cyclopentene-1-yl]hexanoyloxy]propyl]-beta-D-galactopyranose. Library match: name of the MS library used to obtain the candidate molecule annotation.

In addition to this initial selection of metabolites affected by *B. cinerea* infection, we investigated features specifically up-accumulated in low N-grown plants independently of the infection. We hypothesised that these metabolites could be involved in increased basal immunity before any contact with a pathogen. We found 9 features with putative annotations showing this pattern (Figure 5). Four of them, adenylosuccinic acid, unk#12, unk#16 and unk#17, were annotated as N-rich compounds, indicating that they might represent remobilized mobile N sources. Indeed, unk#12, identified as a xanthine, is involved in the purine catabolism pathway, which plays a role in N remobilization whereas adenylosuccinic acid serves as a metabolic intermediate linking amino acid metabolism to both purine synthesis and degradation. Their presence could also contribute to plant defense, for example xanthine is metabolized by xanthine dehydrogenase 1 (XDH1) resulting in ROS production in the epidermal cells, which represses pathogen development, and uric acid production in mesophyll cells, which protects from pathogen induced oxidative stress^48^. Benzoic acid was also more accumulated in mock and infected AWF low N extracts. It could contribute to better resistance as a precursor of salicylic acid or other phenolic plant defense compounds although no depletion was observed after infection at the time point studied.

**Figure 5:**
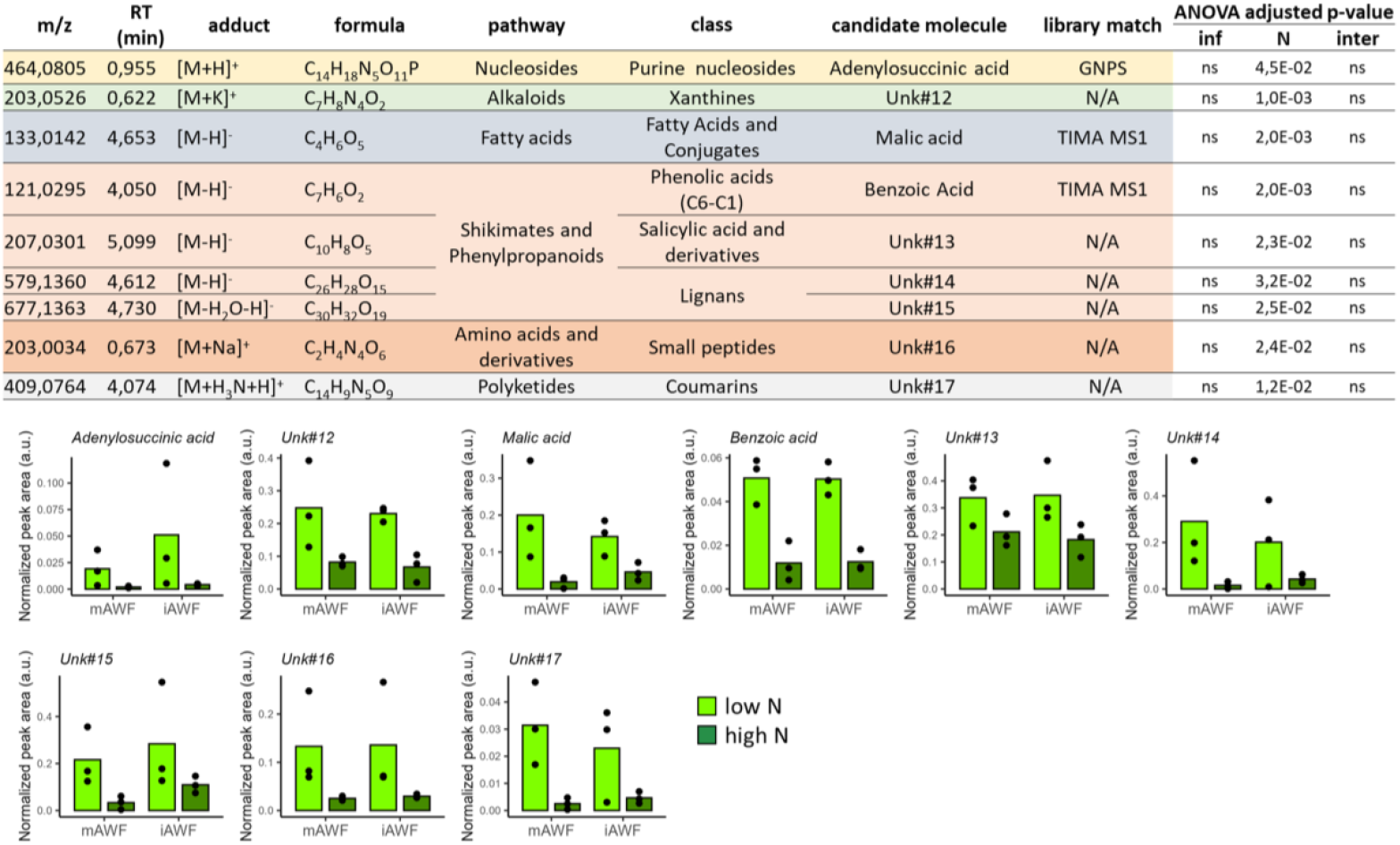
Specialized metabolites enriched in the apoplastic space of plants grown with low nitrogen nutrition. AWF extracts from plants grown in high nitrogen (high N) or low nitrogen (low N) were recovered at 6 h post inoculation with mock (mAWF) or B. cinerea mycelial plugs (iAWF) before analysis with untargeted LC-MS/MS. Tables show for each identified features the detection parameters, the annotation details and the results of a two-way ANOVA for the two factors (inf, N) and their interaction (inter). Barplots represent the mean of normalized peak area (a. u.) for each feature in mAWF and iAWF in low N or high N conditions with each dot representing one independent biological replicate. Library match: name of the MS library used to obtain the candidate molecule annotation.

Thus, in addition to identifying well-characterized compounds involved in defense and in N metabolism, LC-MS/MS analysis of the AWF revealed a new fatty acid conjugate and a new flavonoid compound (Figure 4, unk#8, unk#11) as putative candidates involved in apoplastic defense of plants grown in low N.

### Obtention of quantitative AWF proteomic data

Proteomic analysis was performed on mAWF and iAWF obtained from plants grown in high N and low N conditions in order to examine the changes that occur in the apoplast following *B. cinerea* infection in plants grown under optimal conditions or N nutritional stress. To better understand how the apoplastic space is affected during infection, we added to this dataset iAWF extracts obtained after infection with the *ΔBcacp1* mutant strain of *B. cinerea*, which exhibits reduced pathogenicity^38^.

A set of quantitative proteomic data was obtained from extracted ion chromatogram signals analysis (XIC) in which 913 *A. thaliana* proteins were identified and quantified. GO ontology analysis of the 913 proteins indicated a strong enrichment in different cellular compartments expected for an AWF proteome such as “secretory vesicle”, “apoplast”, “cell wall” and “extracellular region” (Figure 6A). However, some terms related to the chloroplast were also enriched highlighting a partial contamination, as previously described for AWF extracts^19,28^. Moreover, typical functions associated with apoplastic proteins were enriched such as “hydrolase activity”, “peptidase activity” and “peroxidase activity” and the biological processes “mRNA binding”, “cell wall macromolecule catabolic processes”, “detoxification” and “response to stress”. When we looked at the 50 most abundant proteins in the extracts, only six were associated with the chloroplast with Rubisco being the most abundant protein representing 8% of the proteome followed by two apoplastic proteins: GDSL-motif esterase/acyltransferase/lipase 1 (AtGDSL1) and GERMIN-LIKE PROTEIN 1 (AtGLP1), representing respectively around 4 and 3% of the total proteome (Figure 6B). Then, we plotted the relative abundance of all 913 proteins based on their localization and compared it between mAWF, iAWF following infection with the WT strain (iAWF^*B05*.*10*^) and with the mutant strain (iAWF^*ΔBcacp1*^) (Figure 6C). Apoplastic and chloroplastic proteins in mAWF extracts represented respectively 76.4 and 17.3 % of the proteome in low N and 76.9 and 16.6 % in high N which is close to what was observed in Karimi et al. (2025) (Figure 6C). In iAWF^*B05*.*10*^ and iAWF^*ΔBcacp1*^ these proportions where slightly different with an increase of respectively 4.4 and 6.2 % in low N and 4.6 and 6.6% in high N for chloroplastic proteins and a reduction of respectively 4.9 and 6.8 % in low N and 4.7 and 7.2 % in high N for apoplastic proteins. Even though these differences were not always significant (Figure S2), the observed tendency was probably caused by increased cytoplasmic contaminations linked to fungal infection. The rest of the analysis was focused on proteins described as apoplastic or bound to the plasma membrane as they could either be derived from the cleavage of the extracellular domain thus being biologically relevant. We also selected proteins previously identified in proteomic analysis of EVs^16,17^.

**Figure 6:**
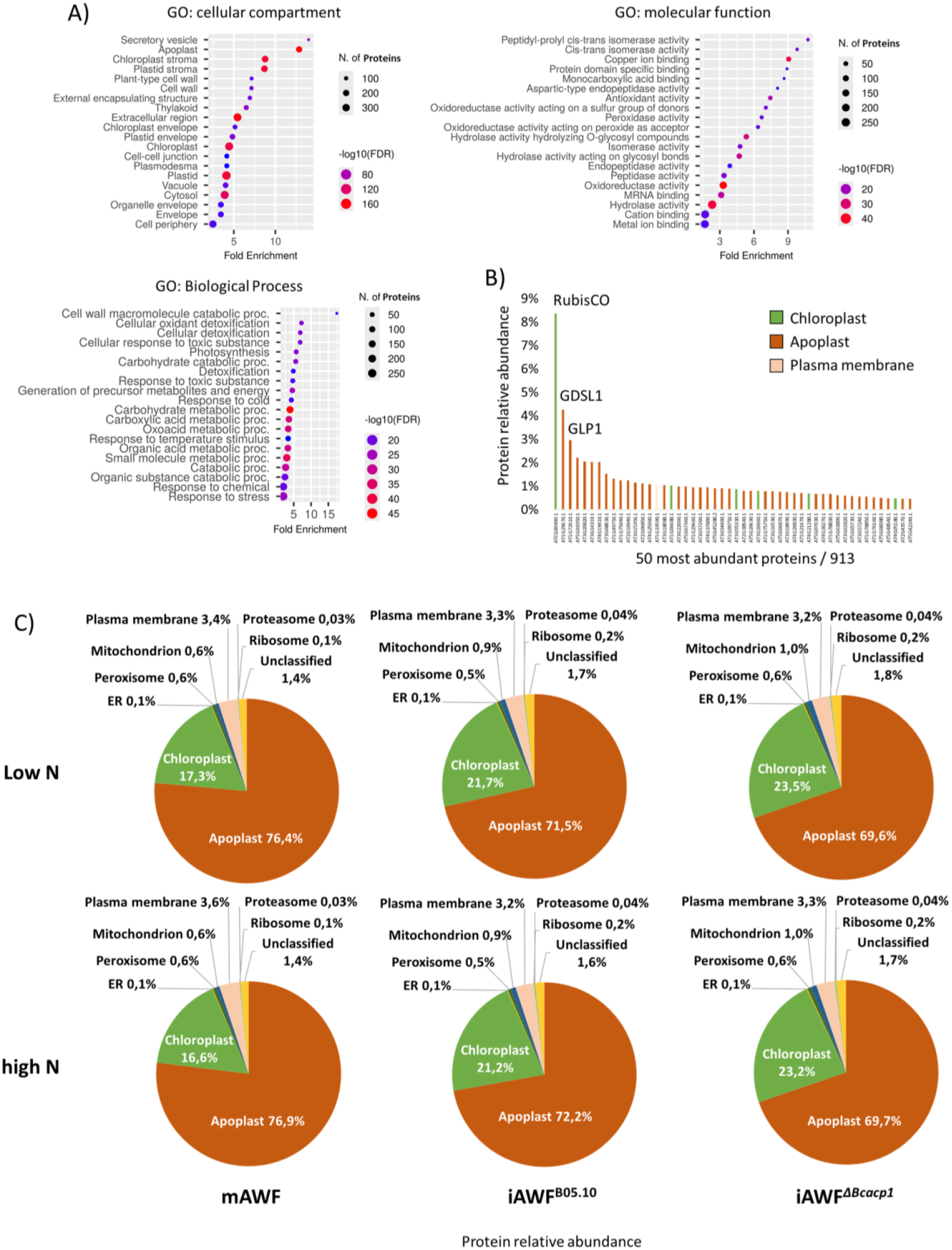
Proteome analysis of AWF extracts reveals a strong enrichment in apoplastic proteins. (A) GO enrichment analysis of identified proteins in AWF extracts. (B) Cellular localization of the 50 most abundant proteins identified in AWF extracts. (C) Relative abundance of proteins based on cellular localization for all three different AWF extracts obtained from plants grown in low N and high N. mAWF: extract obtained from mock-inoculated leaves, iAWF^B05.10^: extract obtained from leaves inoculated with the B05.10 WT strain, iAWF^ΔBcacp1^: extract obtain from leaves inoculated with the ΔBcacp1 mutant strain.

### Changes of known and new proteins involved in defense against B. cinerea in iAWF

Our analysis revealed 120 proteins significantly affected by the infection with the WT *B. cinerea* strain, 36 of which are highly responsive proteins (|log2FC|>1, Table 1). When comparing the accumulation profile of theses protein in the apoplast to the mRNA levels from a previous transcriptomic analysis from detached leaves^38^,19 out of the 36 selected proteins having similar transcriptional patterns (Table 1). In addition, 13 proteins of this list, all more accumulated following infection, were previously detected in EVs that contribute to plant defense^16,17^. Interestingly, a third of these proteins were previously linked to plant defense (underlined in Table 1) and 6 of them, AtDHyPRP1, AtPME17, AtEARLI1, AtPRX62, AtPRX71 and AtPRX21, during *B. cinerea* infection (asterisk in Table 1). Indeed, a higher resistance of the corresponding overexpressor lines and/or increased susceptibility of the corresponding mutants to *B. cinerea* infection was observed^49–57^. Several cell-wall remodelling proteins that differentially accumulated in iAWF remain uncharacterized in the context of defense against the fungus, making them good candidates for future functional studies. Among them, two xyloglucan trans-hydrolases, AtXTH23 and AtXTH32 were respectively up and down accumulated in iAWF samples compared to mAWF and the same trend was found in the RNAseq data. To validate these results, we phenotyped two independent T-DNA insertion lines, *xth32*.*1* and *xth32*.*2*, and found that they were more susceptible to the infection with fungal spores (Figure S3). On the contrary, out of the two independent *xth23* lines we tested, one showed increased resistance to the fungus. Overall, these results highlight the pivotal role of apoplastic proteins in defense against *B. cinerea* and reveal new candidates, such as the remaining 17 highly accumulated proteins, potentially involved in regulating the interaction.

**Table 1:**
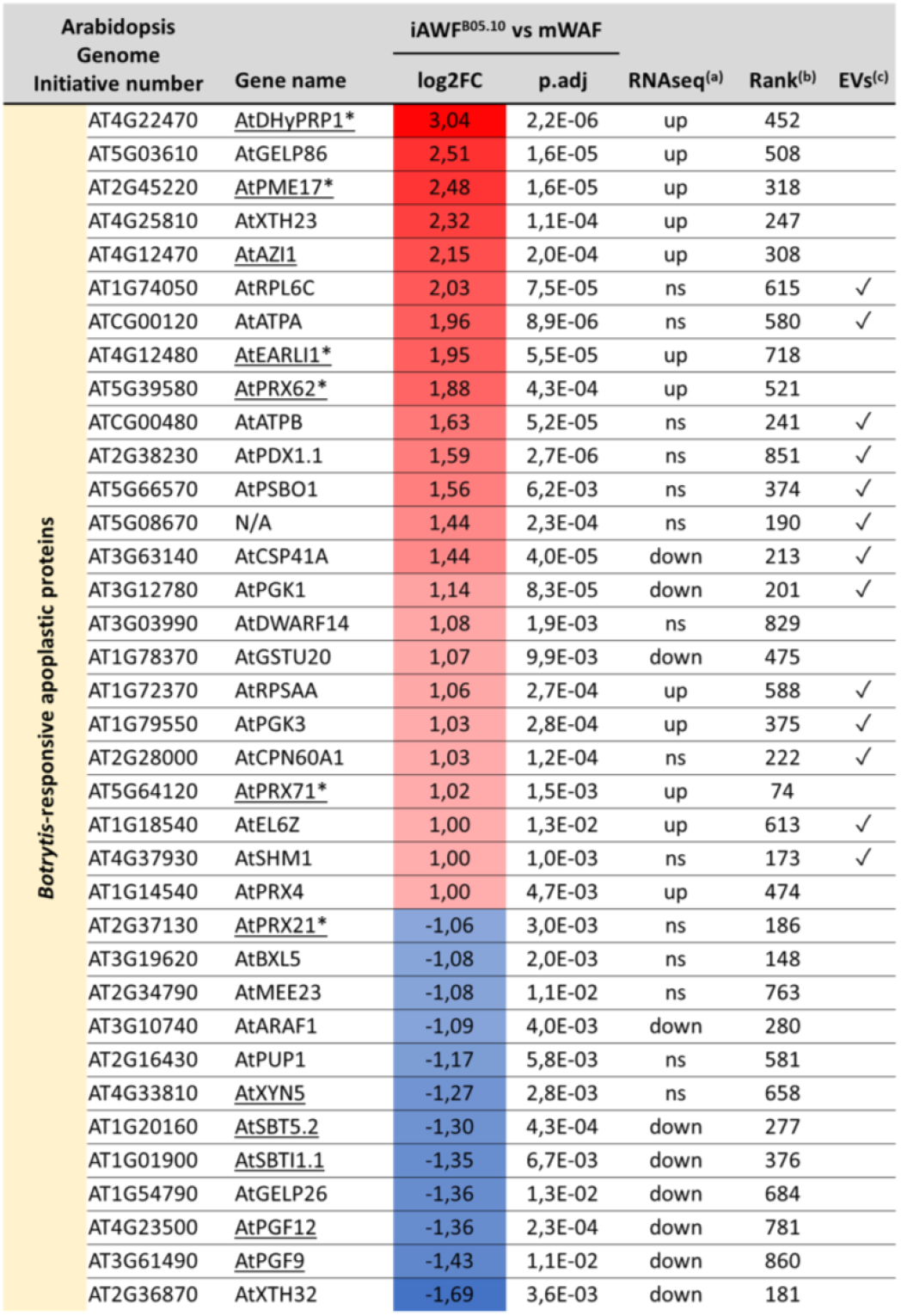
AWF proteins highly responsive to Botrytis cinerea WT strain. ANOVA adjusted p-values <0.05 and |log^2^(FC)|>1. A Tuckey HSD post hoc test was performed to obtain the p.adj values for the comparison mAWF/iAWF^B05.10^. Underline proteins were previously linked to defense against pathogens. *Proteins described as important for B. cinerea resistance. ^(a)^ Regulation of the gene expression in a previously generated RNAseq analysis upon B. cinerea infection of detached Arabidopsis leaves^38. (b)^ Ranking of protein abundance over the 913 detected proteins in AWF extracts. ^(c)^ Proteins detected in proteomic studies focusing on extracellular vesicles^16,17^.

### Effect of the ΔBcacp1 mutation and plant N supply on the AWF proteome

The BcACP1 protease was shown to be transcriptionally up-regulated on infected host plants grown in high N supply and to be an important early-secreted pathogenicity factor^38,58^. To find putative direct plant targets of the BcACP1 fungal protease we looked for proteins up-accumulated in iAWF^*B05*.*10*^ compared to iAWF^*ΔBcacp1*^. However, we could only find five down-accumulated proteins that could be indirect targets of the BcACP1 protease (Table S2). Interestingly two of these proteins were trichome birefringence-like proteins (TBL), including AtTBL38 recently described to be a pectin acetylesterase^59^. Accordingly, cell wall modifications induced in response to the mutant may contribute to its reduced pathogenicity.

Then, we looked for proteins differentially regulated by plant N supply (Figure 7). Seven proteins in the AWF were significantly affected by plant N nutrition with only two of them, AtRAN1 and AtXTH4, also affected by the infection. Only AtRAN1 was more accumulated in AWF from plants grown in low N that are more resistant to the fungus as well as in iAWF compared to mAWF. Interestingly, AtRAN1 was previously described to be involved in plant defense but also to interact with HASTY, which participates in the N starvation response of *A. thaliana*^60–62^. Although AtRAN1 was described to be involved in nuclear transport of different proteins, it has been observed in other AWF-proteomic analysis and detected in EVs^16^. At the transcriptional level, *AtRAN1* was not significantly affected by plant N supply suggesting post-translational regulation of the apoplastic accumulation of AtRAN1 in low N plants.

**Figure 7:**
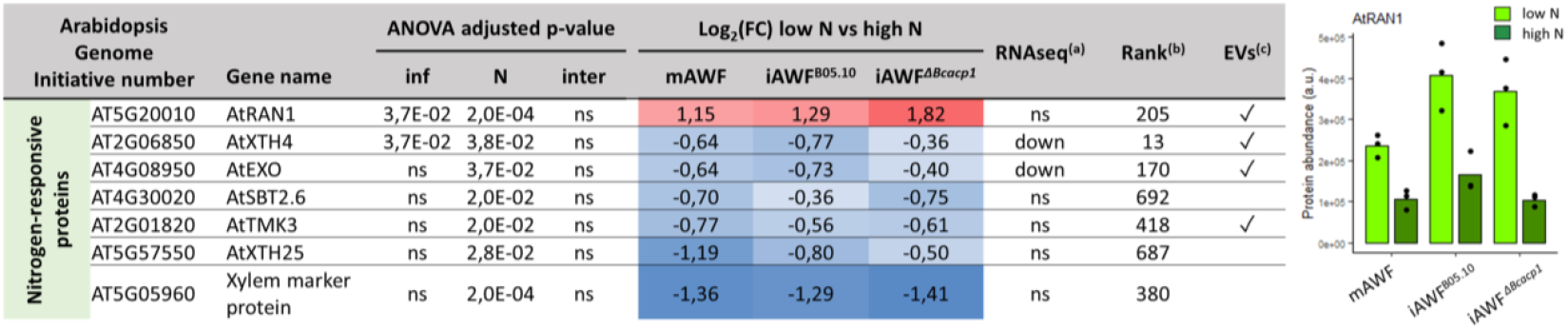
AWF proteins responsive to the plant’s nitrogen supply. The table shows the proteins with an ANOVA adjusted p-values <0.05 for the nitrogen nutrition factor (N). ^(a)^ Regulation of the gene expression in a previously generated RNAseq analysis between plants grown in low N and high^38. (b)^ Ranking of protein abundance over the 913 detected proteins in AWF extracts. ^(c)^ Proteins detected in proteomic studies focusing on extracellular vesicles^16,17^. The histogram shows the protein abundance in all samples for AtRAN1 with the dots representing individual replicates.

### B. cinerea proteins recovered in AWFs extracts

Plant pathogens, particularly fungi, are known to secrete a wide range of proteins during infection. Our proteomic dataset actually contained 68 *B. cinerea* proteins, but these were present in quantities too low to be reliably quantified in the different samples. We therefore focused on the functions of the 37 proteins that were most likely to be secreted, based on their predicted localization or detection in previously described secretomes of the fungus (Table 2).

**Table 2:**
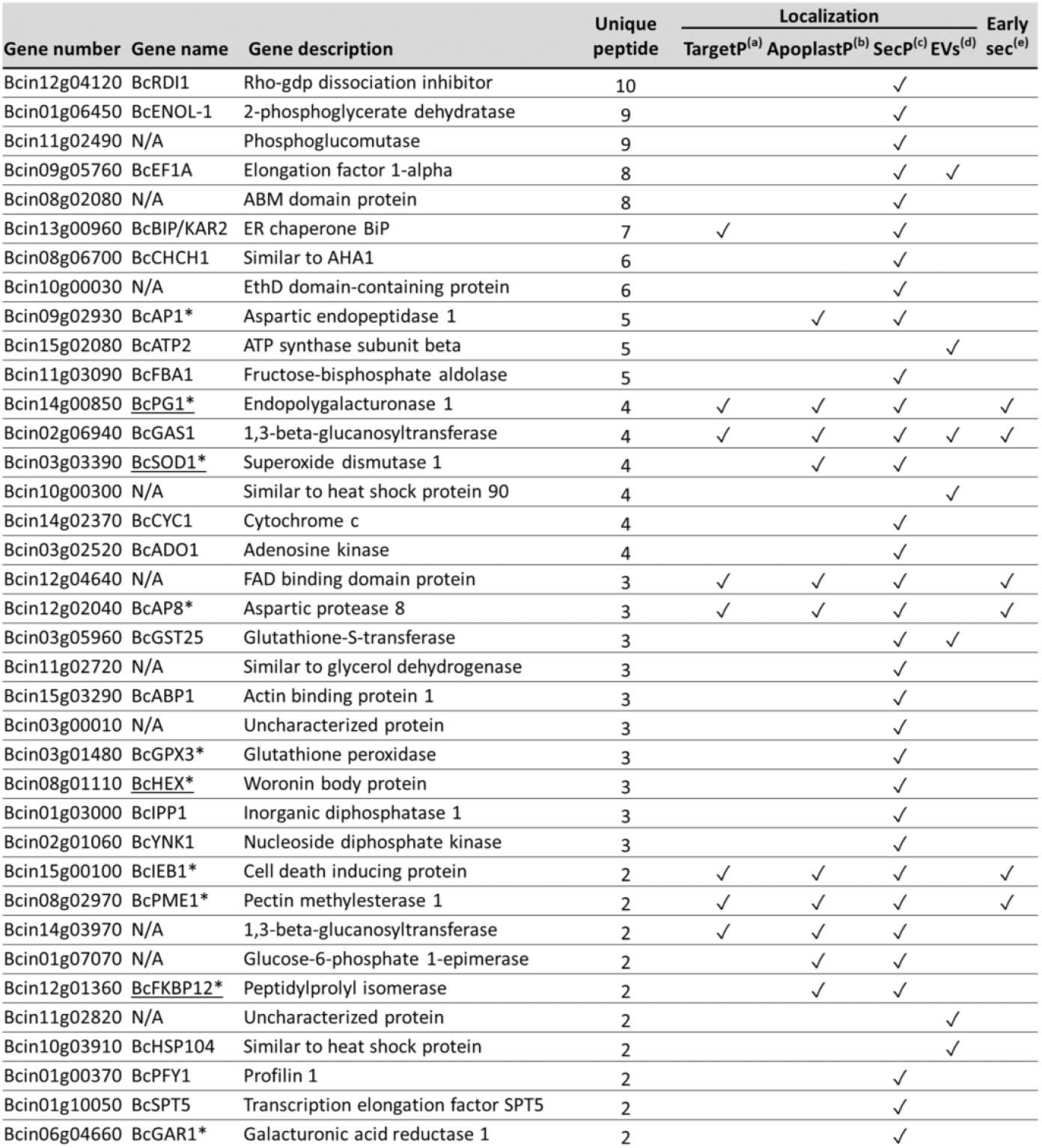
Botrytis cinerea proteins detected in the AWF extracts from infected plants. The table indicates the number of unique peptides detected for each B. cinerea protein, the predicted localization using prediction tools and its detection in other studies. *indicates that a mutant of this gene is published and underlined gene names indicate a role in pathogenicity. ^(a)^Proteins with a signal peptide identified with the TargetP software. ^(b)^ Proteins predicted to be apoplastic using the ApoplastP software. ^(c)^Proteins with a score >0.5 using the SecretomeP software. ^(d)^ Proteins detected in the extracellular vesicle of B. cinerea produced during in vitro growth on cellophane sheets^63. (e)^ Proteins detected in the early secretome of B. cinerea with different plant extracts^64^.

This list of proteins revealed well-known processes of *B. cinerea* pathogenicity: pectin degradation (BcPG1, BcPME1), protein degradation (BcAP8, BcAP1), production of ROS and detoxification (BcSOD1, BcGST25, BcGPX3) and production of specialized metabolites (Bcin10g00030, PKS7 gene cluster). Interestingly, 7 proteins had been previously detected in EVs produced by the fungus during *in vitro* growth^63^, indicating the capture of *B. cinerea* secreted EVs in our iAWF extracts. Six proteins were previously detected by mass spectrometry in the early secretome of *B. cinerea*^*64*^: BcPG1, BcAP8, BcIEB1, BcPME1 and two uncharacterized proteins Bcin12g04640 and Bcin02g06940. By testing the corresponding deletion mutants BcPG1, BcSOD1, BcFKBP12 and BcHEX were previously found involved in pathogenicity^65^ whereas BcIBE1, BcAP8, BcPME1 and BcAP1 were not ^31,66,67^. However, BcIEB1 was shown to promote plant cell death, which is in favour of fungal development and to contribute to pathogenicity in a higher-order mutant^35^. In addition to the fungal proteins already characterized or associated with well-established processes, our proteomic analysis identified 27 proteins predicted to be secreted but not yet studied in the context of infection. Seven of those proteins combined at least two lines of evidence, from prediction tools or other proteomic studies, for being secreted: BcEF1A, BcBIP/KAR2, BcGAS1, Bcin12g04640, BcGST25, Bcin14g03970 and Bcin1g07070. These proteins thus represent promising candidates for future functional characterization.

## Discussion

Recently, increased attention has been given to the apoplast, due to its implication in plant-microbe interactions as well as in senescence, nutrient uptake and response to a large array of stresses^68^. Despite its key role for plant cells, this compartment remains one of the least understood within plant tissues, in part due to its lack of membrane and its difficult extraction process, which can lead to cell damage and contaminations from other compartments. To our knowledge, no study has ever focused on the apoplast fluid (APF) in response to necrotrophic fungi, probably because destruction of plant tissue by the pathogen makes the recovery of clean apoplast washing fluid extracts (AWF) more challenging. In this study, we focused on the early non-destructive phase of *B. cinerea* infection of *A. thaliana* leaves (Figure 1). After checking the low cytoplasmic contamination of our AWF extracts, we could demonstrate experimentally their bioactivity towards *B. cinerea* growth *in vitro* (Figure 2). Interestingly, the growth inhibition observed with mAWF and iAWF was comparable but a higher ROS production by the fungus was detected when incubated with iAWF, suggesting the presence of specific factors in iAWF that induce fungal ROS production. These results clearly indicated the presence of both basal and induced antifungal activity in the leaf apoplast. In order to reveal new differentially regulated proteins and metabolites that are not picked up in optimal conditions, we conjointly studied the immune response with an abiotic stress response by varying plant N nutrition, a factor that influences the susceptibility of *A. thaliana* to *B. cinerea*^10,21,38^.

Many of the molecules affected by *B. cinerea* alone in our multi-omics analysis were already known to be involved in the defense response against this pathogen emphasizing the relevance of focusing on the APF to study plant-pathogen interaction. We found that SA as well as JA and ET, through their precursors ACC and OPDA respectively, were more accumulated in iAWF indicating the ongoing induction of the defense response in the apoplast (Figure 3). In addition, the depletion of metabolites related to energy metabolism or amino acids that we observed is indicative of a shift in resource allocation and a probable pathogen starvation strategy of the host in parallel to direct consumption of nutrients by the fungus. Recently, knocking out UmamiT20 in *A. thaliana*, a plasma membrane-localized broad-spectrum amino-acid transporter, was shown to increase susceptibility to *B. cinerea* infection^9^ illustrating the importance of nutrient fluxes in the APF during the interaction. In parallel, specialized metabolites such as camalexin and some octadecanoids, previously characterized as anti-Botrytis compounds^69^, were strongly accumulated (Figure 4). Moreover, we identified a previously uncharacterized flavonoid compound (unk#11) strongly accumulated in iAWF, particularly in low N. Another example is a fatty acid conjugate compound (unk#8) depleted following infection but only in low N that could be a defense compound precursor. These molecules, which would need structural confirmation, could participate in the defense against *B. cinerea* and the increased resistance of plants grown in low N. Out of 36 highly *Botrytis*-responsive proteins, our comparative proteomic analysis identified 12 proteins described to be involved in plant defense with 6 of them against *B. cinerea* (Table 1). Although the effect was limited, we revealed the contribution of one of the remaining 24 uncharacterized proteins picked up in the analysis, XTH32, in quantitative resistance against the fungus through a genetic approach (Figure S3).

*A. thaliana* plants grown under N limitation (low N) are more resistant to *B. cinerea*^*38*^. We hypothesised that highly accumulated molecules in the mAWF of plants grown in these conditions, regulated or not by the infection itself, could be involved in basal apoplastic immunity against *B. cinerea*. Compared to the strong effect of *B. cinerea* inoculation on the APF composition, the effect of N nutrition was more subtle. Only a few metabolites and one protein showed this pattern, with some previously described to play a role in plant immunity or N metabolism (Figure 5, 7). Interestingly, some metabolites are involved in both N remobilization and defense. For example, xanthine was shown to be involved in *A. thaliana* defense response against powdery mildew^48^. In addition to the metabolites, our proteomic analysis revealed that AtRAN1, a ran GTPase involved in nucleo-cytoplasmique transport^70^, is more accumulated in AWF from plants grown in low N and responds to the fungus (Figure 7). This protein was found to be involved in chitin-induced resistance against *Sclerotinia* s*clerotiorum*, a necrotrophic fungus closely related to *B. cinerea*^62^. Thereby, chitin-triggered defense involving AtRAN1 could be facilitated in low N nutrition conditions leading to a more efficient plant defense induction. Ran GTPases are highly conserved proteins in eukaryotes and were also linked to immunity in mammals^71^. Furthermore, in plants AtRAN1 was shown to be involved in nuclear transport of proteins such as the SA signalling regulator NPR1 but also HASTY involved in the N-starvation response and the production of miRNA^60,72^. However, the presence of AtRAN1 in the apoplast, probably in EVs as observed previously^16,70^, raises questions on its role. It was shown in mammalian cells that this class of Ran GTPases can be secreted in exosome-type EVs, enabling the intercellular transport of nuclear proteins from one cell to the other^70^. Since AtRAN1 protein accumulation in the APF is unlinked to mRNA levels, post-transcriptional regulation of AtRAN1 secretion in EVs might explain the data. Further study is needed to determine if the regulation of AtRAN1 secretion in EVs and its interaction with specific nuclear cargoes are involved in the plant multi-stress response to *B. cinerea* and N limitation. Future studies using proximity labelling techniques coupled with APF proteomics may reveal the nature of AtRAN1 nuclear cargo(es) transported via EVs during the interaction.

Finally, we detected *in vivo* early secreted proteins of *B. cinerea* in the APF. Well known pathogenicity factors of the fungus were detected such as the very abundant BcPG1 polygalacturonase. Seven out of the 37 *B. cinerea* selected proteins in the APF were previously detected in vesicle during *in vitro* growth^63^ highlighting the presence of EVs of fungal origin in our extracts. However, 27 detected proteins in this work still need to be characterized in the context of the interaction. Based on prediction tools and recent proteomic studies, seven of these proteins show a high likelihood of being secreted. Their presence in the APF battlefield during the early and decisive phase of the fungal infection make them promising candidates for further functional studies.

Overall, our study reveals the potential of studying the apoplastic compartment during plant-pathogen interactions (Figure 8) to discover new players, even in a well-described pathosystem. By studying the apoplastic space in the early phase of the interaction between *A. thaliana* and *B. cinerea*, we were able to identify new putative plant defense proteins and metabolites as well as pathogen effectors. In the future, it could be valuable to search for peptides present in this compartment from both plant and fungal origin using peptidomics since the understanding on these molecular players is currently still limited.

**Figure 8:**
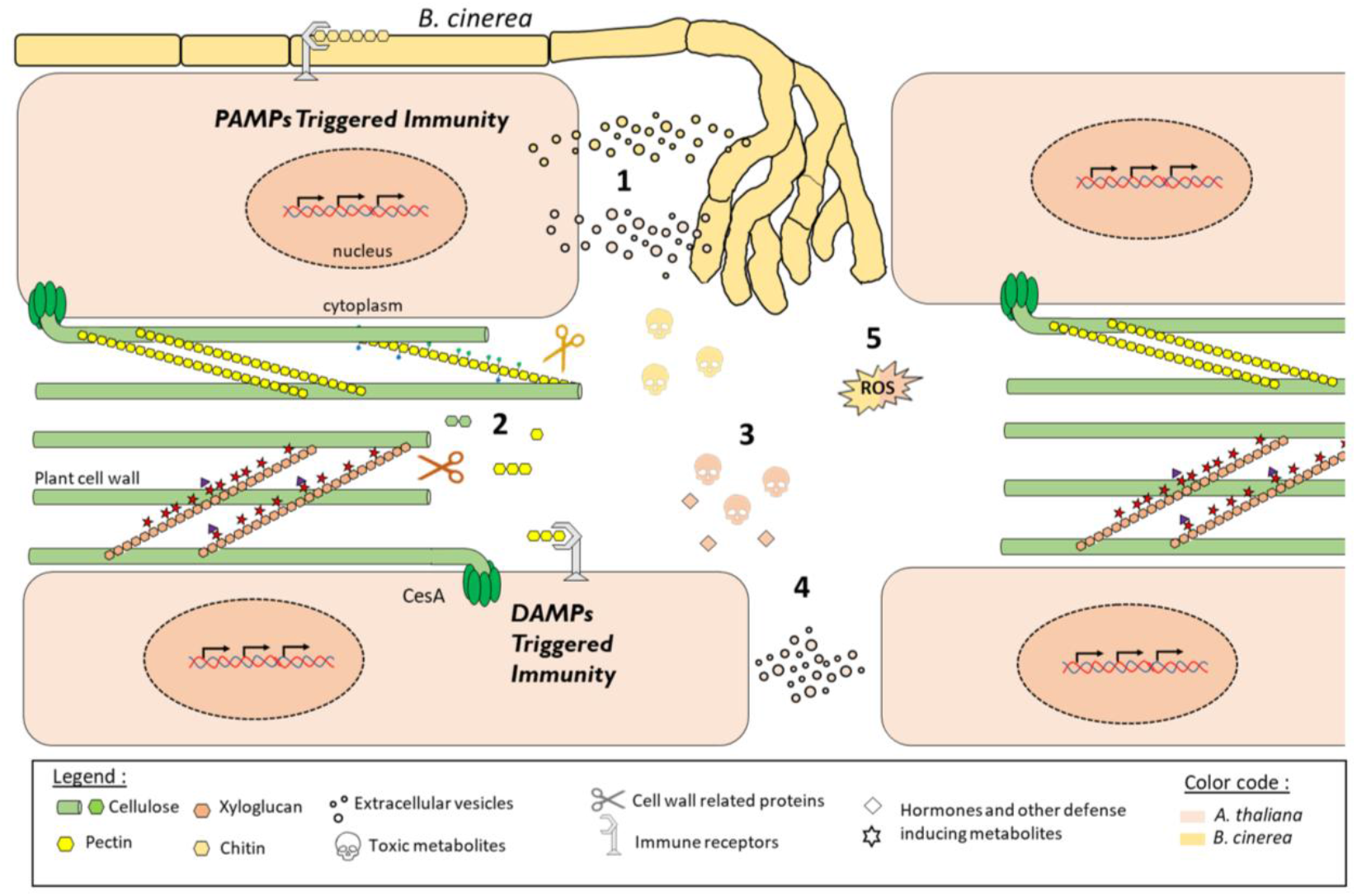
Important processes taking place in the apoplast during plant-pathogen interaction identified by multi-omics profiling of the apoplastic fluid. 1) Pathogen and host proteins loaded in extracellular vesicles (EVs). 2) Cell wall remodelling through accumulation and depletion of host proteins (AtPME17, AtXTH32 and others) as well as accumulation of oligomers of cell wall polysaccharides (galacturonic acid, cellobiose, rhamnose) from the activity of pathogen CWDEs (BcPG1 and BcPME1). 3) Production of host toxins (camalexin), specialized metabolites (oxylipins, flavonoids) and hormones (OPDA, ACC and SA), building the defense response. 4) Cell-cell communication through EVs carrying nuclear proteins transporters (AtRAN1) propagating the immune response to neighbouring cells. 5) Production of reactive oxygen species (ROS), secretion of redox enzymes and antioxidant compounds from the pathogen (BcSOD1 and others) and the host (AtPRX71 and others).

## Materials and Methods

### Fungal and plant material

*B. cinerea* strain B05.10 collected from *Vitis* in Germany^73^ was used throughout this work as the wild-type (WT) reference strain. *B. cinerea* was routinely cultured at 23°C under constant light on potato dextrose agar (PDA, Difco) to collect spores or on malt agar to obtain mycelial plugs (MA, 1% malt extract, 1.2% agar). The *ΔBcacp1* mutant strain of the fungus was kindly received from Pr. Nathalie Poussereau (Université Lyon 1). *A. thaliana* ecotype *Col-0* was obtained from the INRAE-Versailles collection whereas T-DNA insertion mutants *xth32*.*1* and *xth32*.*2* as well as *xth23*.*1* and *xth23*.*2* were obtained from the SALK collection. For AWF extraction, plants were cultured in growth chambers (65% relative humidity, 8 hr photoperiod, 21°C) for 6 weeks on nonsterile sand and were supplied with a nutrient solution containing either 10 mM or 0.5 mM NO_3_^-^. For phenotyping of the plant mutant lines, plants were cultured for 2 weeks on nonsterile sand supplied with a nutrient solution containing 5 mM NO_3_^-^.

### Infection experiments

For AWF extraction, the fungus was cultured for four days on MA agar. Rosettes of 6-week-old *A. thaliana* plants were cut at their base and a mycelial plug (1 cm diameter), taken from the actively growing edge, was inverted onto the upper surface of each mature leaf of the rosette to maximize the interacting surface. Rosettes were then kept in large square dishes under high humidity conditions at 23°C and under continuous light for 6 h before AWF extraction. For qPCR analysis phenotyping of the plant mutant lines, 2-weeks-old *A. thaliana* plants were inoculated in the green house with drops of 2 µL containing 0.5.10^6^ spores.ml^-1^ on each leaf. Lesions were measured using ImageJ at 2 dpi.

### Detection of ROS and cell death

For coloration assays, plants were infected as described above and leaves were cut from rosettes 6 hpi or 24 hpi prior to coloration. Mycelial plugs were gently removed from the 6 hpi collected leaves but kept on the 24 hpi ones because of their fragility. Detection of H2O2 production was achieved by 3,3’-diaminobenzidine (DAB) staining. Briefly, samples were immerged in an aqueous solution containing 1 mg/mL DAB (pH 3.6, Sigma) and incubated overnight at room temperature on a rotary shaker. To de-stain the leaves, chlorophyll was extracted by incubation in 75% ethanol. To detect cell death, Lactophenol Trypan Blue (TB, Sigma) was performed as described in^74^. Pictures were taken on a ZEISS Axio Zoom V16. For oxidative stress revelation after *in vitro* growth of *B. cinerea* on 96-wells plates, the plates were centrifuged for 10 min at 1500 g and the supernatant removed. Then, each well was filled with 300µL of the same above-mentioned DAB solution. Pictures of the plates were taken after 4 h incubation with DAB.

### Apoplastic Washing Fluid (AWF) extraction

AWF extraction was performed with the infiltration-centrifugation method as described previously^21^. At 6 hpi, leaves were cut from rosettes and mycelial plugs were removed. After a brief wash in ddH2O (double distilled water), leaves were immersed in the infiltrating solution containing either ddH2O for metabolomic analysis and nephelometry or 0.3 M Mannitol with 0.2 M CaCl2 pH 6.9 for proteome analysis^40^. Infiltration was performed in vacuum pump. Several pressure cycles of -25 Pa were applied until all the leaves were infiltrated. The infiltrated leaves were then gently dried with paper and inserted into a 20mL syringe. The syringe is then placed inside a 50 mL conical tube. Collection of AWF samples was achieved by centrifugation of the syringe placed in a conical tube at 1000g for 20 min at 4°C. The resulting AWF was centrifuged again for 5 min at 15000 g at 4°C to remove cellular debris, the recovered supernatant collected corresponded to the final extract. The protein content of each extract was then measured by the Bradford method using BSA as standard. AWF extracts were assayed for malate dehydrogenase (MDH) activity to detect cytoplasmic contaminations. MDH was assayed at room temperature as followed: 20µL of extracts were added to reaction mixture containing a buffer (HEPES 100mM pH7), 10 % glycerol, 150 µM NADH and 2 mM oxaloacetate. The oxidation of NADH was followed at 340_nm_. The AWF dilution factor was also assayed for ddH_2_O infiltrated extract as described in O’Leary et al (2014). In each condition some leaves were infiltrated with 50 µM indigo carmine (IC) in ddH_2_O and the absorbance at 610 nm (OD_610_) of the resulting AWF extracts was measured and compared to control conditions in order to calculate the dilution factor: (OD_610 IC AWF_ - OD_610 IC_) / ((OD_610 IC AWF_ - OD_610 IC_)-(OD_610 H2O AWF - OD610 H2O_)).

### Test for antifungal effect of ddH2O AWF extracts

Growth rates were measured by using a nephelometer with the procedure developed by Joubert et al (2010). For inoculum preparation, conidia were collected from 8-day-old on solid medium cultures and suspended in Potato Dextrose Broth (PDB) to obtain a concentration of 10^4^ conidia/mL. For each condition, three wells of a 96 wells-microplate were filled with 300 µL of suspension, supplemented with 30µL of ddH2O AWF extracts. Four different AWF extracts from different extractions were tested. Growth was automatically recorded each 15 minutes during 48h (193 cycles) using a nephelometric reader (NEPHELOstar Plus, BMG Labtech, Omega 5.70), equipped with a 635-nm laser as radiation source. Measurements were done with a laser beam focus of 2.5 mm and an intensity of 80%. The 96-well plates were shacked at 400 rpm, with a double orbital movement between readings. Data were exported from NephelOstar Plus software in ASCII format and further processed in Microsoft Excel. For each condition and each timepoint, a Relative Nephelometric Unit (RNU) value was calculated as the mean of the eight measurements. For each condition, the slope was calculated during the exponential phase using measurements between 20 hpi and 48 hpi to estimate the growth rate. The experiment was done three times.

### AWF primary metabolites content analysis

Metabolomic analysis of primary metabolites with GC-MS was performed as described previously ^75–77^. For the extraction, 50 µL of AWF were diluted with 100 µl of frozen (-20°C) water:acetonitrile:isopropanol (2:3:3) containing ribitol at 4 µg/ml and extracted for 10 min at 4°C with shaking at 1400 rpm in an Eppendorf Thermomixer. Insoluble material was removed by centrifugation at 20000g for 5 min. A blank tube underwent the same steps as the samples. Supernatant was dried overnight at 35 °C in a Speed-Vac vacuum centrifuge. For derivatization, 10 µl of 20 mg/ml methoxyamine in pyridine were added to the samples and the reaction was performed for 90 min at 28°C under continuous shaking in an Eppendorf thermomixer. 90µl of N-methyl-N-trimethylsilyl-trifluoroacetamide (MSTFA) (Aldrich 394866-10x1ml) were then added and the reaction continued for 30 min at 38°C. After cooling, 45 µl were transferred to an Agilent vial for injection. Injections started 4 hours after the end of derivatization. The whole series was first injected in splitless mode and then in split mode (1/30). Four different standards mix were injected at the beginning and the end of the analysis as well as three independent derivatizations of the quality control (beginning, middle and end) for monitoring the derivatization stability. Samples were randomized. An alkane mix (C10, C12, C15, C19, C22, C28, C32, C36) was injected in the middle of the queue for external RI calibration. Injection volume was 1 µl. The instrument was an Agilent 7890A gas chromatograph coupled to an Agilent 5975C mass spectrometer. The column was a Rxi-5SilMS from Restek (30 m with 10 m integraguard column). The liner (Restek # 20994) was changed before each derivatization series. Oven temperature ramp was 70 °C for 7 min then 10 °C/min to 330 °C for 5 min (run length 38 min). Helium constant flow was 0.7 mL/min. Temperatures were the following: injector: 250°C, transfer line: 290°C, source: 250 °C and quadripole 150 °C. 5 scans per second were acquired spanning a 50 to 600 Da range. Instrument was tuned with PFTBA with the 69 m/z and 219 m/z of equal intensities. 5 scans per second were acquired. The split mode conditions were: 70°C for 2 min then 30°C per min to 330 °C for 5 min. Helium constant flow was 1 mL/min.

For data processing, Raw Agilent datafiles were converted in NetCDF format and analyzed with AMDIS (http://chemdata.nist.gov/mass-spc/amdis/). An home retention indices/ mass spectra library built from the NIST, Golm (http://gmd.mpimp-golm.mpg.de/), and Fiehn databases and standard compounds was used for metabolites identification. Peak areas were also determined with the Targetlynx software (Waters) after conversion of the NetCDF file in masslynx format as well as TargetSearch. AMDIS, Target Lynx and TargetSearch in splitless and split 30 modes data were compiled in one single Excel File for comparison. After blank mean substraction peak areas were normalized to ribitol and rresh weight. For absolute quantification, a response coefficient was determined for 4 ng each of a set of 103 metabolites, respectively to the same amount of ribitol. This factor was used to give an estimation of the absolute concentration of the metabolite in what we may call a “one-point calibration”. Metabolites rich in nitrogen (basic aminoacids and polyamines) gave several analytes (up to 5 for glutamine and asparagine). The peak area as total ion current equivalent of these analytes were summed to express the contents of these metabolites.

### AWF specialized metabolites content analysis

Concerning the specialized metabolites, we performed a UPLC-MS/MS analysis. Data were acquired using a UHPLC system (Ultimate 3000 Thermo) coupled to quadrupole time of flight mass spectrometer (Q-Tof Impact II Bruker Daltonics, Bremen, Germany). A Nucleoshell RP 18 plus reversed-phase column (2 x 100 mm, 2.7 μm; Macherey-Nagel) was used for chromatographic separation. The mobile phases used for the chromatographic separation were (A) 0.1% formic acid in H2O and (B) 0.1% formic acid in acetonitrile. The flow rate was of 400 μL/min and the following gradient was used: 95% of A for 1-min, followed by a linear gradient from 95% A to 80% A from 1 to 3-min, then a linear gradient from 80% A to 75% A from 3 to 8-min, a linear gradient from 75% A to 40% A from 8 to 20-min. 0% of A was hold until 24-min, followed by a linear gradient from 0% A to 95% A from 24 to 27-min. Finally, the column was washed by 30% A for 3.5min then re-equilibrated for 3.5-min (35-min total run time). Data-dependent acquisition (DDA) methods were used for mass spectrometer data in positive and negative ESI modes using the following parameters: capillary voltage, 4.5kV; nebuliser gas flow, 2.1 bar; dry gas flow, 6 L/min; drying gas in the heated electrospray source temperature, 140ºC. Samples were analysed at 8Hz with a mass range of 100 to 1500 m/z. Stepping acquisition parameters were created to improve the fragmentation profile with a collision RF from 200 to 700 Vpp, a transfer time from 20 to 70 µs and collision energy from 20 to 40 eV. Each cycle included a MS fullscan and 5 MS/MS CID on the 5 main ions of the previous MS spectrum.

For data processing, the .d data files (Bruker Daltonics, Bremen, Germany) were converted to .mzXML format using the MSConvert software (ProteoWizard package 3.0^78^) mzXML data processing, mass detection, chromatogram building, deconvolution, samples alignment, and data export were performed using MZmine 2.52 software (http://mzmine.github.io/) for both positive and negative data files. The ADAP chromatogram builder method^79^ was used with a minimum group size of scans of 3, a group intensity threshold of 1000, a minimum highest intensity of 1500 and m/z tolerance of 10 ppm. Deconvolution was performed with the ADAP wavelets algorithm using the following setting: S/N threshold 8, peak duration range = 0.01-1 min RT wavelet range 0.02-0.2 min, MS2 scan were paired using a m/z tolerance range of 0.01 Da and RT tolerance of 0.1 min. Then, isotopic peak grouper algorithm was used with a m/z tolerance of 10 ppm and RT tolerance of 0.1. All the peaks were filtered using feature list row filter keeping only peaks with MS2 scan. The alignment of samples was performed using the join aligner with an m/z tolerance of 10 ppm, a weight for m/z and RT at 1, a retention time tolerance of 0.1 min. A first research in library with Mzmine was done with identification module and “custom database search” to begin the annotation with our library, currently containing 78 annotations (RT and m/z) in positive mode and 38 in negative mode, with RT tolerance of 0.3 min and m/z tolerance of 0.0025 Da or 6 ppm.

Metabolites annotation of the total dataset was performed in two consecutive steps. First, the obtained RT and m/z data of each feature were compared with our home-made library containing more than 150 standards or experimental common features (RT, m/z). Second, the ESI- and ESI+ metabolomic data used for molecular network analyses were searched against the available MS2 spectral libraries (Massbank NA, GNPS Public Spectral Library, NIST14 Tandem, NIH Natural Product), with absolute m/z tolerance of 0.02, 4 minimum matched peaks and minimal cosine score of 0.65. Then, for selected interesting metabolites presented in Figure 4 and 5, Sirius software (https://bio.informatik.uni-jena.de/software/sirius/) was used. Sirius is based on machine learning techniques that used available chemical structures and MS/MS data from chemical databanks to propose structures of unknown compounds^80–82^. Searches for analogues based on molecular spectra and structures were performed using MS2Query (https://github.com/iomega/ms2query; de Jonge et al. 2023) and in order to optimise annotation confidence by focusing exclusively on the species taxon, we used the Taxonomically Informed Metabolite Annotation (TIMA; https://github.com/taxonomicallyinformedannotation/tima) tool for the *Arabidopsis thaliana* structure-organism pair.

### Proteome analysis of AWF extracts

The LC-MS/MS analyses have been done at the PAPPSO proteomics platform (pappso.inra.fr/). Following short 1D protein electrophoresis, in-gel digestion was performed using standard in gel digestion as described in Recorbet et al (2021). After drying, sample were reconstituted in 30 µl of 2% acetonitrile (ACN) 0.1 Formic acid in water. Sample were diluted to 50 ng per µl of peptide. Sample were analyzed using a nanoElute (Bruker, Billerika USA) coupled to timsTOF Pro (Bruker). 2 µl of sample were loaded on a Acclaim PepMap C18 Trap Column, (100 Å, 5 µm, 100 µm X 20 mm, ThermoScientific) with 0.1% HCOOH (v/v) in water for 1 min at 300 bar. Peptides were further separated, on a Aurora ultimate CSI C18 column (120 Å, 1,7 µm, 75 µm X 250 mm, ionopticks). The mobile phase consisted of a gradient of solvents A: 0.1% HCOOH (v/v), in water and B: 100% ACN (v/v), 0.1% HCOOH (v/v) in water. Separation was set at a flow rate of 0.25 μl/min using a linear gradient of solvent B from 2 to 13% in 42 min, followed by an increase to 20% in 23 min and an increase to 30% in 5 min. Finally, Buffer B was increased to 85% in 5 min and stay at this value for 12 min. Nebulization was performed using a Captive spray interface at 1.6 kV applied on liquid junction. The timsTOF Pro acquisition have been realized with timsControl (version 3.0.21) with the following settings: Mass Range 100 to 1700 m/z, 1/K0 Start 0.7 V ⋅s/cm2 End 1.10 V ⋅s/cm2, Ramp time 180 ms, Lock Duty Cycle to 100%, Dry Gas 3 l/min, Dry Temp 180°C in PASEF mode. PASEF settings: 10 MS/MS scans (total cycle time 1.3 sec), charge range 2-5, active exclusion for 0.4 min, Scheduling Target intensity 14000, Intensity threshold 1000 and CID collision energy from 20 to 59 eV. Database search was performed against araport 11 (https://www.arabidopsis.org/download/indexauto.jsp?dir=%2Fdownload_files%2FProteins%2FArap_ort11_protein_lists), Botrytis cinerea database (https://www.uniprot.org/uniprotkb?query=(taxonomy_id:999810) 16338 entries) and a custom contaminant database using X!Tandem^84^ (see Table S3 for details). Data filtering and protein inference was perform using i2MassChroQ^85^. A protein was considered validify its Evalue was less than 0.00001 and if it was identified by at least two distinct peptides, each with an Evalue less than 0.01. These filters produced a FDR of 0.15 at the peptide-spectrum match level, 0.31 at the peptide level and 0 at the protein level. Protein quantification was perform using MassChroQ^86^ as described in Balliau et al (2018). For XIC analysis, only peptide presenting less than 10% of missing value, with a retention time standard deviation after alignment lesser than 20s and presenting a correlation between each peptide for a given protein higher than 0,5 was used for protein quantity computation. Protein quantity was computed as the sum of peptides intensity after imputation of missing value (method irmi from VIM Package) on non-shared peptides. The mass spectrometry proteomics data have been deposited to the ProteomeXchange Consortium via the PRIDE^88^ partner repository with the dataset identifier PXD073646.

### Statistical analysis

At least three independent biological replicates of apoplastic extracts were analysed in each omic experiment and to look for antifungal activity. Infection experiments were repeated at least three times with more than 15 leaves in each condition. Statistical analysis was performed with the R studio software. After a log10 transformation of the quatitative values for proteins and metabolites, a two-way ANOVA test was performed. When needed, a tukey’s HSD post hoc test for factor levels comparison was performed following the ANOVA for omics experiments or a two-sample t-test for other experiments.

## Acknowledgment

Proteomics analyses were performed on the PAPPSO platform (http://pappso.inrae.fr) which is supported by Paris Saclay University (https://www.universite-paris-saclay.fr/en), INRAE (http://www.inrae.fr), CNRS (http://www.cnrs.fr), AgroParisTech (https://www.agroparistech.fr/en), the Ile-de-France regional council (https://www.iledefrance.fr/education-recherche), IBiSA (https://www.ibisa.net), Saclay Plant Sciences-SPS (ANR-17-EUR-0007), and Plant2Pro (https://plant2pro.fr/). A.D was supported by a doctoral fellowship from U. Paris Saclay. This work has benefited from the support of IJPB’s Plant Observatory platforms PO-Plants and P0-Chem. The IJPB benefits from the support of Saclay Plant Sciences-SPS (ANR-17-EUR-0007). This work was supported by INRAE-BAP (BcEqolisin). We thank Célia Perdrizet, Thomas Tempestini and François Pélé for their help with the analysis of the *A. thaliana* mutants.

