## Supplemental Data for "Apoplast multi-omics profiling during fungal infection uncovers new players of basal and early induced immunity"

***Table S1:*** *Comparison of qualitative markers of the Apoplastic washing fluids (AWF) extraction between AWF extracts obtained from mock inoculated plants (mAWF) and inoculated with the B. cinerea WT strain (iAWF).*

| | | mAWF | iAWF |
|---|---|---|---|
| **Mannitol-$CaCl_2$ infiltration** | Activité MDH ($U.ml^{-1}$) | 0.018 +/- 0.0035 | 0.024 +/- 0.003 |
| | Protein concentration ($\mu g.\mu L^{-1}$) | 0.26 +/- 0.05 | 0.35 +/- 0.1 |
| **dd$H_2$0 infiltration** | Activité MDH ($U.ml^{-1}$) | 0.0072 +/- 0.006 | 0.01 +/- 0.0025 |
| | Protein concentration ($\mu g.\mu L^{-1}$) | 0.190 +/- 0.041 | 0.193 +/- 0.023 |
| | AWF dilution factor | 1.67 +/- 0.38 | 1.65 +/- 0.35 |

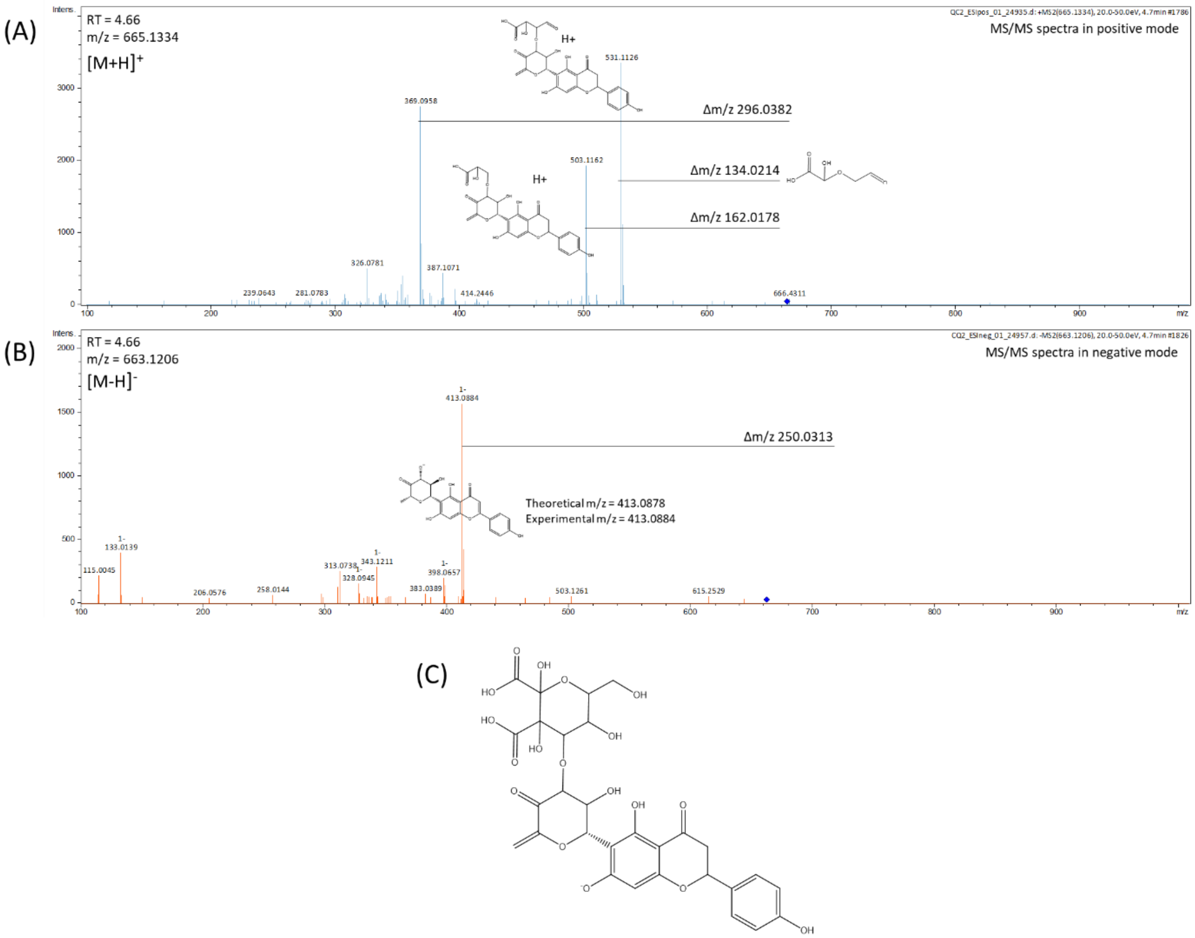


***Figure S1: Identification of the new aglycone flavonoid accumulated in conditions of plant resistance to the fungal infection.*** *MS/MS fragmentation spectra in positive (A) and negative mode (B) showing the putative fragments of the molecule corresponding to the proposed structure (C).*

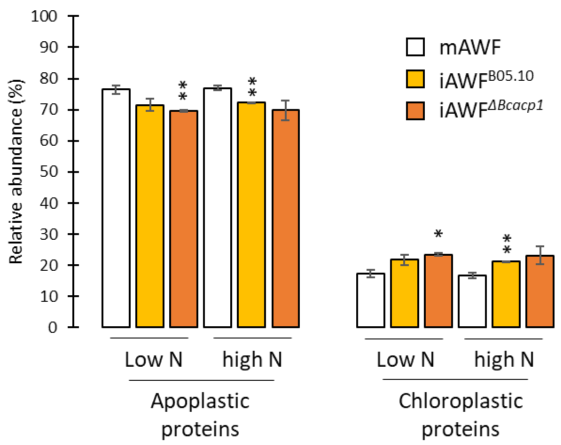


***Figure S2: Proteome analysis of AWF extracts reveals a strong enrichment in apoplastic proteins.*** *Relative abundance of proteins based on cellular localization for all three different AWF extracts obtained from plants grown in low N and high N. mAWF: extract obtained from mock- inoculated leaves, iAWF$^{B05.10}$: extract obtained from leaves inoculated with the B05.10 WT strain, iAWF$^{\Delta Bcacp1}$: extract obtain from leaves inoculated with the ΔBcacp1 mutant strain. A two sample t-test was performed to compare mAWF to the iAWF extracts in each N condition: *p<0.05, **p<0.01.*

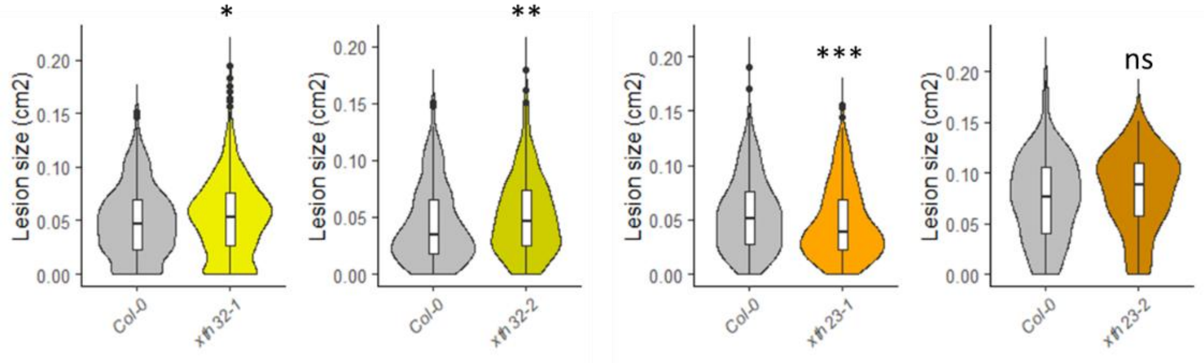


***Figure S3: Role of the apoplastic proteins XTH32 and XTH23 in plant defense against B. cinerea.*** *Plot of the lesions on A. thaliana detached leaves at 2 dpi with the WT strain of the fungus B05.10 inoculated on col-0, the two xth32 and the two xth23 mutant lines. A two sample t-test was performed to compare the lesions between col-0 and the mutant lines: ns for not significant, *p<0.05, **p<0.01, ***p<0.001.*

***Table S2: AWF proteins responsive to the ΔBcacp1 mutant of B. cinerea**. A tuckey HSD post hoc test was performed to obtain the p.adj values. [(b)]Ranking of protein abundance over the 913 detected proteins in AWF extracts.[(c)]Protein detected in proteomic studies focusing on extracellular vesicles[16,17].*

| | Arabidopsis Genome Initiative number | Gene name | iAWF$^{B05.10}$ vs mWAF | | iAWF$^{\Delta Bcacp1}$ vs mWAF | | iAWF$^{\Delta Bcacp1}$ vs iAWFB05.10 | | Rank[(b)] | EVs[(c)] |
|---|---|---|---|---|---|---|---|---|---|---|
| | | | $log_2$(FC) | p.adj | $log_2$(FC) | p.adj | $log_2$(FC) | p.adj | | |
| ***Bcacp1*-responsive proteins** | AT3G04720 | AtPR4 | 0.31 | ns | -0.88 | 1.1E-02 | -1.19 | 3.0E-03 | 493 | |
| | AT3G14850 | AtTBL41 | -0.80 | ns | -1.77 | 3.7E-03 | -0.96 | 4.9E-02 | 727 | |
| | AT1G29050 | AtTBL38 | -0.11 | ns | -0.71 | 1.8E-03 | -0.61 | 6.9E-03 | 372 | |
| | AT5G66590 | AtCAP61 | -0.50 | 9.8E-03 | -0.90 | 8.1E-05 | -0.41 | 3.2E-02 | 338 | |
| | AT4G38670 | thaumatin | -0.49 | 6.4E-03 | -0.82 | 6.4E-05 | -0.33 | 3.6E-02 | 494 | |

*Table S3: Identification parameter for protein identification from A. thaliana and B. cinerea protein databases.*

### Protein identification

#### Xtandem Run information

| param | value |
|---|---|
| version | X! Tandem Piledriver (2015.04.01.1) |
| sequence source #1 | Araport11_genes.201606.pep.fasta |
| sequence source #2 | contaminants_standarts.fasta |
| sequence source #3 | uniprotKB_B_cinerea_220329.fasta |

#### Spectra filtering

| param | value |
|---|---|
| parent monoisotopic mass error minus | 50 |
| parent monoisotopic mass error plus | 50 |
| parent monoisotopic mass error units | ppm |
| parent monoisotopic mass isotope error | yes |
| maximum parent charge | 5 |
| fragment mass type | monoisotopic |
| fragment monoisotopic mass error | 50 |
| fragment monoisotopic mass error units | ppm |
| total peaks | 100 |
| neutral loss mass | 18.01057 |
| neutral loss window | 0.05 |
| dynamic range | 100.0 |
| minimum fragment mz | 150.0 |
| minimum parent m+h | 500.0 |
| minimum peaks | 15 |

#### Enzymatic cleavage

| param | value |
|---|---|
| cleavage site | [RK]\|{P} |
| cleavage semi | no |
| quick pyrolidone | yes |
| stP bias | yes |
| quick acetyl | yes |

**Amino acid modification**

| param | value |
|---|---|
| modification mass | 57.02146@C |
| potential modification mass | 15.99491@M |

**Refinement module**

| param | value |
|---|---|
| use refine | yes |
| maximum valid expectation value | 0.01 |
| spectrum synthesis | yes |
| unanticipated cleavage | no |
| modification mass | 57.02146@C |
| point mutations | no |
| potential modification mass | 15.99491@M |
| use potential modifications for full refinement | yes |
| cleavage semi | yes |
| spectrum synthesis | yes |

**Scoring**

| param | value |
|---|---|
| minimum ion count | 4 |
| maximum missed cleavage sites | 1 |
| cyclic permutation | yes |
| include reverse | yes |
| b ions | yes |
| y ions | yes |
